# Multidimensional *T*_1_-*T*_2_ Relaxation Imaging *In Vivo* on a Portable 0.064 T MRI Scanner

**DOI:** 10.64898/2026.09.04.749459

**Authors:** Ella Wilczynski, Kulam Najmudeen Magdoom, Alexandru V Avram, Nathan H Williamson, Dan Benjamini, Silvina G Horovitz, Peter J Basser

## Abstract

**Purpose:** To demonstrate the feasibility of whole-brain multidimensional *T*_1_-*T*_2_ relaxation imaging on a portable 0.064 T MRI scanner.

**Methods:** A two-dimensional inversion-recovery, fast spin-echo acquisition was used to jointly encode *T*_1_ and *T*_2_ relaxation. Joint relaxation distributions were reconstructed voxel-wise using marginal-distribution constrained optimization (MADCO). The approach was evaluated in a quantitative relaxation MRI phantom and subsequently applied *in vivo* in a healthy volunteer. Joint distributions, marginal distributions, distribution-derived measures, and statistical dependencies between the two relaxation dimensions were examined.

**Results:** Phantom joint relaxation estimates showed good overall agreement with mono-exponential relaxation measurements and previously reported values at 0.064 T. The *in vivo* acquisition provided whole-brain coverage over a broad range of relaxation weightings and tissue contrasts. Reconstructed joint *T*_1_-*T*_2_ distributions showed spatially organized features across the relaxation space, including broad and overlapping relaxation components that were not fully represented by single-value relaxation maps or by either 1D marginal distribution alone. Selected regions of the joint relaxation space produced component maps with distinct spatial patterns. Statistical analysis further showed a dependence between the two relaxation dimensions, supporting the presence of added information in the joint distribution that is lost when treating *T*_1_ and *T*_2_ separately.

**Conclusion:** Whole-brain multidimensional *T*_1_-*T*_2_ relaxation imaging is clinically feasible on a portable 0.064 T MRI scanner. Joint relaxation distributions provide information beyond single-value relaxation mapping and may support further developments of quantitative multidimensional imaging at ultra-low field.

## Introduction

Portable ultra-low-field (ULF) MRI scanners (i.e., operating below 0.1 T) are emerging as accessible alternatives to conventional MRI scanners. They have lower infrastructure requirements and cost and can operate in point-of-care and resource-limited environments (1; 2; 3). These permanent-magnet scanners do not require cryogenic cooling or conventional shielded MRI infrastructure and have been used for neuroimaging in intensive-care, emergency, mobile, and geographically distributed settings (4; 5; 6). Recent studies have also demonstrated their potential for evaluating a range of intracranial pathologies (7; 8). These advantages may be particularly important where access to conventional MRI remains limited. For example, the Ultra-low-field Neuroimaging In The Young (UNITY) initiative is deploying 0.064 T MRI across multiple sites in low- and middle-income countries (LMICs) to study early brain development (9). As access to portable ULF MRI expands, there is also a need to extend the quantitative information available from these scanners beyond structural imaging alone.

Quantitative MRI (qMRI) goes beyond anatomical imaging by estimating tissue properties such as *T*_1_ and *T*_2_, which can provide information related to tissue composition and microstructure. A range of approaches have been developed for quantitative relaxation mapping, including conventional relaxometry and more highly encoded methods such as MR Fingerprinting (10; 11). Because relaxation times vary with magnetic field strength, measurements obtained at conventional field strengths do not necessarily translate directly to ULF. Relaxation properties therefore need to be characterized specifically at these lower field strengths (12). At ULF, previous studies have demonstrated *in vivo* quantitative relaxation measurements in both adult (13; 14; 15) and neonatal populations (16). More recent work has also demonstrated repeatable low-field *T*_1_ mapping across sites and portable MRI scanners using accelerated acquisition and reconstruction approaches (17). These studies establish the feasibility and value of quantitative relaxation mapping at ULF, however, conventional relaxation maps typically represent each voxel by a single estimated *T*_1_ or *T*_2_ value. Such scalar measures cannot describe the heterogeneous relaxation environments that may coexist within a voxel.

Rather than estimating a single voxel-averaged value, relaxation spectroscopic MRI aims to characterize the distribution of MR-sensitive parameters within each voxel, including *T*_2_ (18), *T*_1_ (19), and mean diffusivity (MD) (20). These distributions can reveal multiple relaxation or microstructural environments that are not represented by a single scalar measurement. Multidimensional MRI (mdMRI) extends this concept by jointly encoding multiple MR-sensitive parameters and reconstructing their joint distribution within each voxel (21; 22). In particular, joint *T*_1_-*T*_2_ distributions describe how relaxation components are related across the two dimensions. This relationship is lost when *T*_1_ and *T*_2_ are considered independently (23). Multidimensional approaches have demonstrated sensitivity to tissue microstructure and compartmental composition (24; 25; 26; 27; 28; 29; 21), and have been applied to investigate placental dysfunction (30), astrogliosis (31), diffuse axonal injury (32), and diffusivity-anisotropy domains *in vivo* (33; 34; 35).

Translating multidimensional MRI to portable ULF scanners presents additional technical challenges. Multidimensional relaxation imaging requires sampling a large jointly encoded parameter space, resulting in long acquisition times and increased susceptibility to noise, motion artifacts, and reconstruction instability. These challenges are amplified at ULF by reduced signal-to-noise ratio (SNR), limited gradient performance, and practical constraints on scan duration. Although multidimensional imaging has been demonstrated at conventional field strengths, its application to portable ULF scanners remains largely unexplored. To our knowledge, voxel-wise joint *T*_1_-*T*_2_ relaxation distributions have not previously been reported *in vivo* on a portable ULF MRI scanner.

In this work, we present a multidimensional *T*_1_-*T*_2_ relaxation imaging framework for a portable 0.064 T MRI scanner using an inversion-recovery fast spin-echo (IR-FSE) acquisition. Joint *T*_1_-*T*_2_ distributions were reconstructed on a voxel-wise basis using marginal-distribution constrained optimization (MADCO) (36; 21) and evaluated first in a quantitative relaxation MRI phantom and subsequently *in vivo* in a healthy volunteer. We also examined *T*_1_-*T*_2_ distribution-derived measures, spatial patterns within the relaxation spectrum, and statistical dependence between the two relaxation dimensions. This proof-of-concept study establishes the feasibility of whole-brain multidimensional relaxation imaging at ULF and provides a basis for future studies addressing specific biological and clinical questions.

## Theory

### Multidimensional Relaxation Modeling

Multidimensional nuclear magnetic resonance (NMR) is well established in chemical spectroscopy, porous media NMR, and structural biology (37). Its translation to MRI has been more difficult because multidimensional relaxometry requires large amounts of data and long scan times. Approaches such as compressed sensing (38) and MADCO (39) have substantially reduced these data requirements. These developments have made voxel-wise multidimensional relaxation measurements more practical while maintaining spectral accuracy and precision.

The relaxation-weighted signal acquired in each voxel, *S*(*TI, TE, TR*), can be modeled as an integral over the joint distribution of the longitudinal and transverse magnetic relaxation rates, 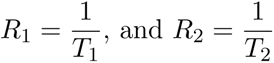, respectively, weighted by their corresponding relaxation kernels:

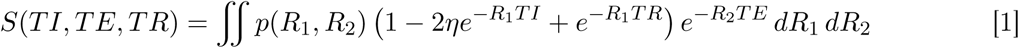

The parameter *η* represents the apparent inversion efficiency. The form of Eq. 1 resembles a multidimensional Laplace transform. In practice, the joint distribution is obtained by treating the problem as a Fredholm integral equation of the first kind, which is discretized and solved by applying additional constraints (40).

For discrete relaxation rates *R*_1_*_,n_* and *R*_2_*_,n_*, let *S_m_* = *S*(*TI_m_, TE_m_, TR_m_*) denote the measured signal for the *m^th^* combination of *TI*, *TE*, and *TR*. The corresponding signal predicted by the discretized model is *S*_0_(Ψ*p*)*_m_*:

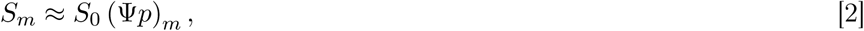

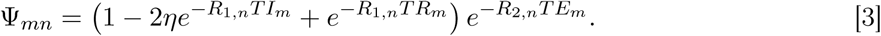

Here, Ψ*_mn_* is the *m* × *n* encoding kernel, and *p* is the discretized joint probability distribution whose *n^th^* element, *p_n_*, corresponds to the relaxation-rate pair (*R*_1_*_,n_, R*_2_*_,n_*). The approximate equality accounts for the difference between the measured and predicted signals, which is minimized in the constrained optimization framework described below.

### Constrained Estimation of the Joint Relaxation Distribution

Estimating multidimensional relaxation distributions is an ill-posed inverse problem, so some form of regularization and additional constraints are required. In MADCO, the solution is limited to positive relaxation rates with *R*_1_ *< R*_2_. The joint distribution, *p*(*R*_1_*, R*_2_), must also be nonnegative and normalized to one. Independently estimated one-dimensional marginal distributions further restrict the admissible joint *R*_1_-*R*_2_ parameter space (39).

The estimation of the joint distribution, *p*(*R*_1_*, R*_2_), can be formulated as the constrained convex optimization problem

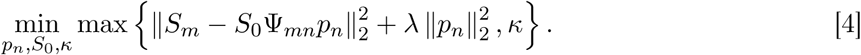

In Eq. 4, the first term measures the difference between the measured and predicted signals. The *ℓ*_2_ term regularizes the joint distribution, with its contribution controlled by *λ*. The optimization is subject to the following physical, probabilistic, and marginal-distribution constraints:

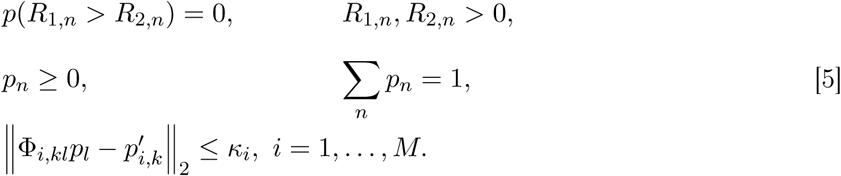

The first four constraints define a physically admissible, nonnegative, normalized joint probability distribution over the 2D relaxation-rate space. The final constraint requires agreement with the independently estimated marginal distributions within a tolerance *κ_i_*. Here, Φ*_i,kl_* maps the joint distribution to its *i^th^*marginal, and *p_i,k_*’ is the corresponding independently estimated marginal.

## Methods

All experiments were performed on portable 0.064 T MRI scanners (Hyperfine SWOOP^®^, Guilford, CT, USA). A research agreement with the manufacturer allowed us to modify the pulse sequence timing and implement multidimensional contrast encoding. We used a two-dimensional IR-FSE acquisition to jointly encode longitudinal and transverse relaxation. The acquisition was repeated over multiple inversion time (*TI*) and repetition time (*TR*) combinations, with multiple echo times (*TE*) sampled within each FSE echo train. These relaxation-weighted images were then used to estimate the joint *T*_1_-*T*_2_ distributions. The acquisition framework is shown in Figure 1.

**Figure 1:**
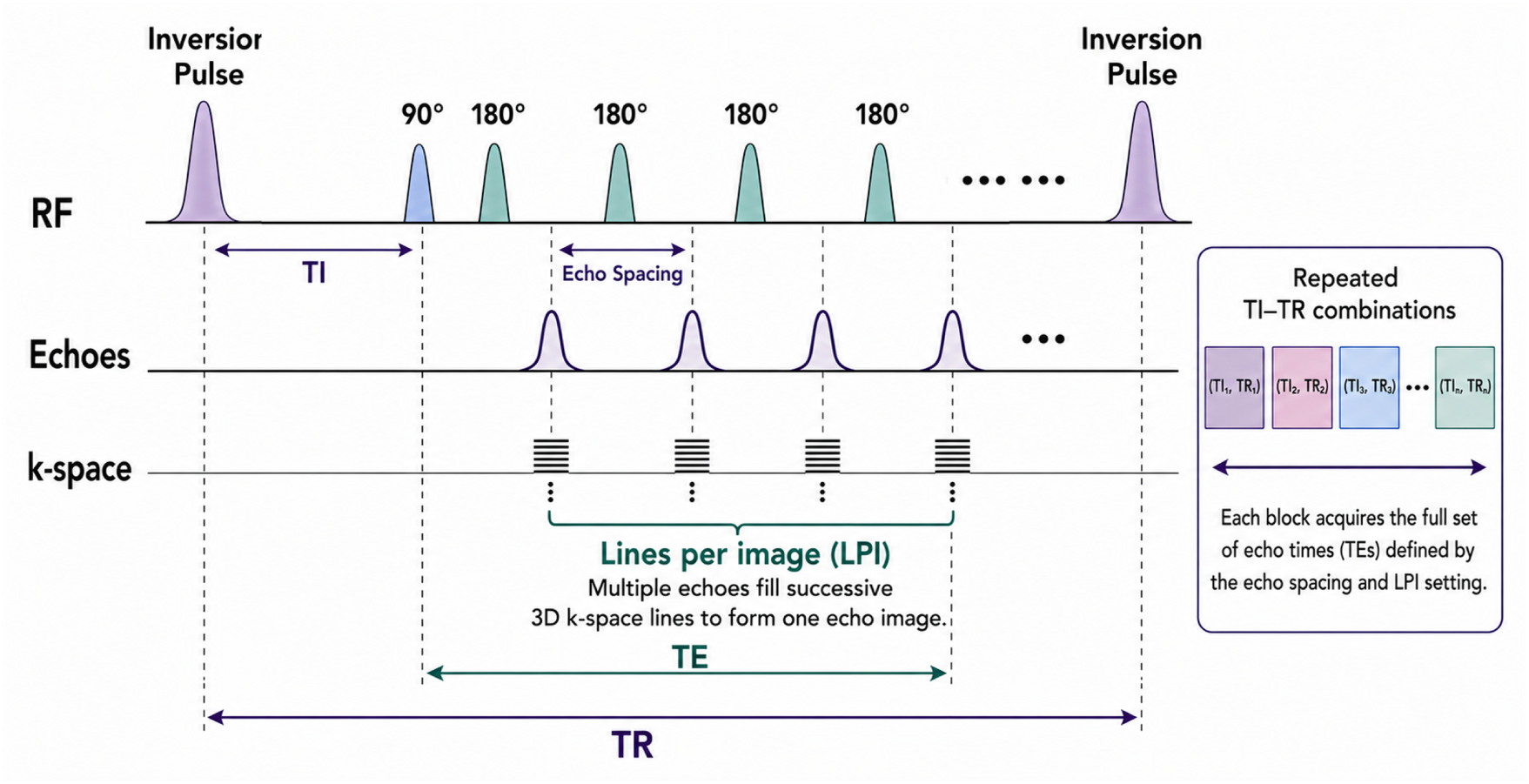
IR-FSE pulse sequence and multidimensional relaxation-encoding strategy. An adiabatic inversion pulse is followed after inversion time (*TI*) by a 90^◦^ excitation pulse and a train of 180^◦^ refocusing pulses separated by the echo spacing. Multiple echoes are acquired within each repetition, with each echo contributing one line in 3D k-space. The Lines Per Image (LPI) parameter determines the number of echoes contributing to each reconstructed echo image. The effective echo time (*TE*) is the interval between the 90^◦^ excitation pulse and the final echo contributing to a given echo image. Repetition time (*TR*) is defined between successive inversion pulses. The sequence block is repeated for multiple *TI* and *TR* combinations, and the full set of echo times is acquired for each combination to provide joint *T*_1_-*T*_2_ relaxation encoding.

### Phantom Study

#### Data Acquisition

Phantom data were acquired using a commercially available quantitative MRI phantom (Model 137, CaliberMRI, Boulder, CO, USA), hereafter referred to as the UNITY phantom (41). The phantom contains tissue-mimicking compartments with varying concentrations of NiCl_2_ and MnCl_2_, covering a broad range of *T*_1_ and *T*_2_ values. It also includes an array of polyvinylpyrrolidone (PVP) compartments for diffusion calibration, geometric fiducials, and other quality-control features. A detailed description of the phantom design and characterization has been reported previously (41).

Data were acquired on a Hyperfine SWOOP portable 0.064 T MRI scanner (hardware version 1.8, software version 9.0.1). We used the two-dimensional IR-FSE acquisition described above and shown in Figure 1. The protocol included 32 inversion times ranging from 22 to 5000 ms and 80 echo images ranging from approximately 6.5 to 522 ms. The repetition time (*TR*) was 5700 ms. Images were acquired with an in-plane resolution of 2.0 × 2.0 mm^2^, a slice thickness of 4.17 mm, and a total acquisition time of approximately one week. A complete summary of the acquisition protocol is provided in Table 1.

**Table 1:**
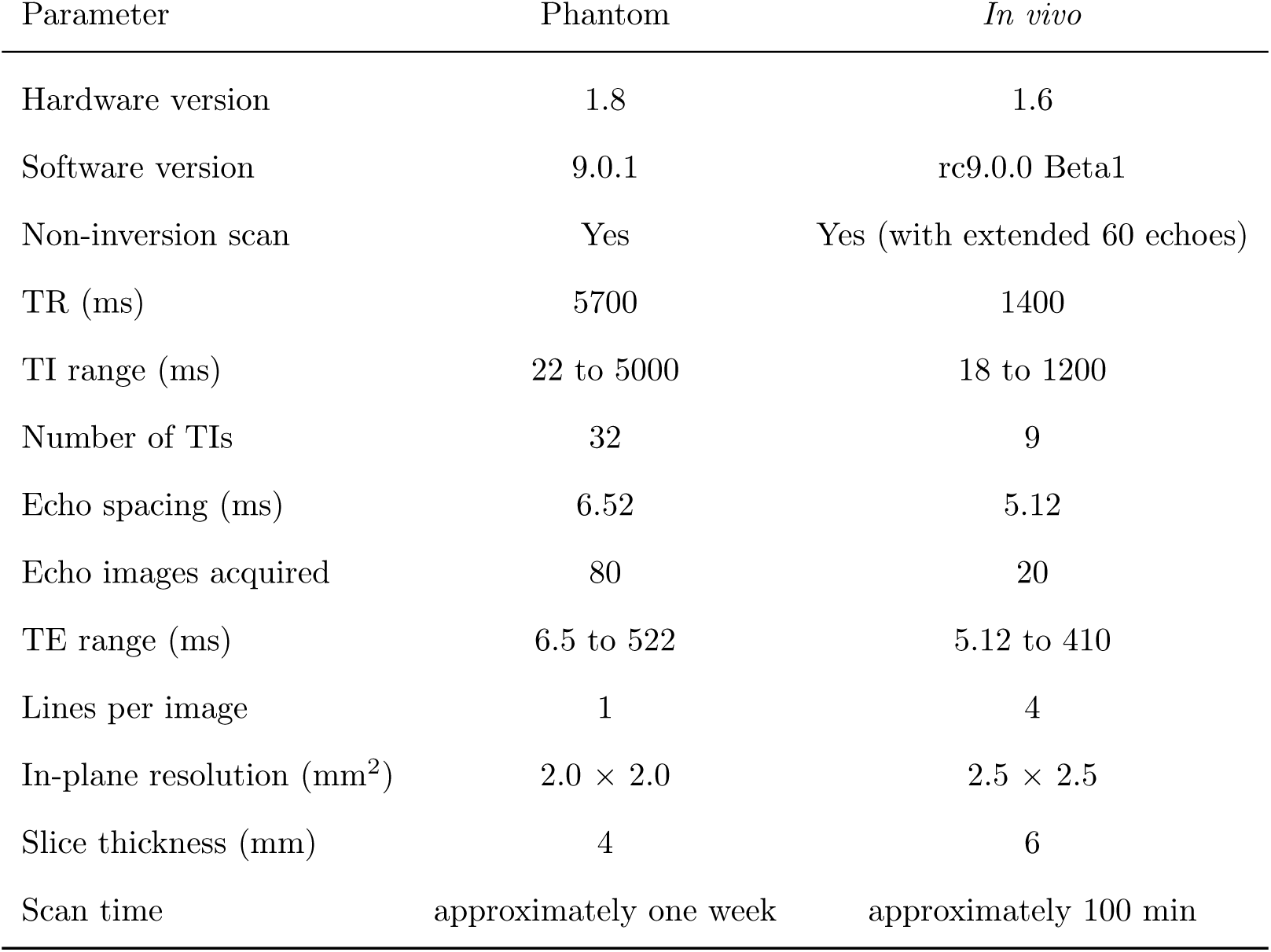
Acquisition parameters for the phantom and *in vivo* multidimensional IR-FSE protocols.

| Parameter | Phantom | *In vivo* |
| --- | --- | --- |
| Hardware version | 1.8 | 1.6 |
| Software version | 9.0.1 | rc9.0.0 Beta1 |
| Non-inversion scan | Yes | Yes (with extended 60 echoes) |
| TR (ms) | 5700 | 1400 |
| TI range (ms) | 22 to 5000 | 18 to 1200 |
| Number of TIs | 32 | 9 |
| Echo spacing (ms) | 6.52 | 5.12 |
| Echo images acquired | 80 | 20 |
| TE range (ms) | 6.5 to 522 | 5.12 to 410 |
| Lines per image | 1 | 4 |
| In-plane resolution (mm<sup>2</sup>) | $2.0 \times 2.0$ | $2.5 \times 2.5$ |
| Slice thickness (mm) | 4 | 6 |
| Scan time | approximately one week | approximately 100 min |

#### Data Analysis

Phantom data were analyzed using mono-exponential and joint relaxation fitting. Each phantom compartment was designed to be chemically homogeneous, so we assumed a single relaxation component. *T*_1_ and *T*_2_ were estimated independently from the inversion recovery and multi-echo datasets using voxel-wise mono-exponential least-squares fitting:

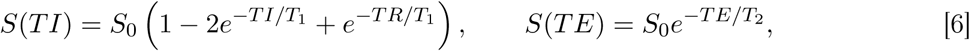

where *S*_0_ represents the equilibrium signal amplitude. Only voxel-wise fits with a coefficient of determination *R*^2^ *>* 0.95 were retained for subsequent analysis.

Joint relaxation fitting used the complete multidimensional phantom dataset and the discrete forward model in Eq. 2, with the encoding kernel defined in Eq. 3. Under the single-component assumption, the joint relaxation-rate distribution for each compartment was modeled as a delta function, *p*(*R*_1_, *R*_2_) = *δ*(*R*_1_ – *R*_1_’)*δ*(*R*_2_ – *R*_2_’). The relaxation times are then *T*_1_ = 1/*R*_1_’ and *T*_2_ = 1/*R*_2_’, and the multidimensional signal model reduces to:

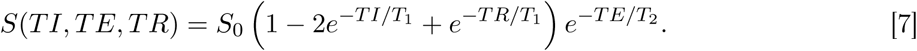

Goodness of fit for the mono-exponential *T*_1_ and *T*_2_ estimates obtained using Eqs. 6 was assessed using linear regression, coefficient of determination (*R*^2^), and root-mean-square error (RMSE). For each phantom compartment, we calculated mean *T*_1_ and *T*_2_ values and their standard errors using regions of interest based on the phantom geometry.

Quantitative validation of the joint *T*_1_-*T*_2_ analysis focused on phantom compartments with relaxation properties spanning the range relevant to human brain tissue. We compared the joint relaxation estimates with the mono-exponential *T*_1_ and *T*_2_ estimates. Where available, we also compared them with previously published MRI and NMR measurements obtained at 0.064 T (41; 42). Agreement between the estimated and reference relaxation times was evaluated relative to the line of identity.

### *In Vivo* Human Brain Imaging

#### Data Acquisition

*In vivo* MRI experiments were performed in one healthy adult volunteer (28-year-old male) under an Institutional Review Board (IRB)-approved research protocol (NCT06203626). Data were acquired on a Hyperfine SWOOP portable 0.064 T MRI system (hardware version 1.6, software version rc9.0.0_Beta1). We used the two-dimensional IR-FSE acquisition described above and shown in Figure 1.

The volunteer was scanned using nine inversion times ranging from 18 to 1200 ms. An additional acquisition without an inversion pulse was used to sample near-fully relaxed magnetization. *TR* was fixed at 1400 ms for all inversion times. Four lines per image (LPI = 4) were acquired. Multiple echoes within each FSE echo train therefore filled successive lines in 3D k-space to form an echo image, as shown in Figure 1. The echo spacing was 5.12 ms, with *TE* values extending to 409.6 ms. Images were acquired with an in-plane resolution of 2.5 × 2.5 mm^2^, a slice thickness of 5.8 mm, and a field of view (FOV) of 180 × 220 × 156 mm^3^. Complete acquisition parameters are provided in Table 1. The total acquisition time was approximately 100 minutes.

#### Data Analysis

Prior to spectral reconstruction, the MR images were processed in the following order: 1) random noise was suppressed using a Marchenko-Pastur principal component analysis (MP-PCA) algorithm (43) implemented in the DIPY software environment (44); 2) inter-scan subject motion, odd-even echo errors, and eddy-current-induced geometric distortions were reduced by registering all volumes to the first echo acquisition using a 3D affine transformation (45; 46) implemented in the FSL software environment (47); and 3) signal polarity was restored for magnitude images using the algorithm described in (48) to account for the inversion.

Mono-exponential *T*_1_ and *T*_2_ maps were generated using the signal models defined in Eq. 6. *T*_1_ was estimated from the minimum-*T E* inversion-recovery images and *T*_2_ from the multi-echo non-inversion acquisition. Representative white matter, gray matter, and cerebrospinal fluid (CSF) voxels were selected for comparison with previously reported relaxation values.

Voxel-wise joint relaxation distributions were reconstructed from the *in vivo* dataset using MADCO and the *ℓ*_2_-regularized optimization and constraints defined in Eqs. 4 and 5. The joint distribution, *p*(*R*_1_*, R*_2_), was represented on a logarithmically spaced 21 × 21 grid of relaxation rates, corresponding to *T*_1_ and *T*_2_ ranges of approximately 18-5,600 ms and 10-1,000 ms, respectively. The solution was required to be nonnegative and normalized, with the marginal distributions restricting the admissible *R*_1_-*R*_2_ parameter space according to Eq. 5. This space included regions encompassing 95% of the marginal probability, with adaptive gridding used to increase sampling density where needed and reduce it elsewhere. The regularization parameter *λ* was determined independently in each voxel using the *S*-curve method. The apparent inversion efficiency, *η*, was estimated voxel-wise and incorporated into the longitudinal relaxation kernel used for the joint reconstruction (Figure 6b).

For visualization and comparison with mono-exponential measurements, the reconstructed spectra were converted to relaxation-time coordinates using *T*_1_ = 1*/R*_1_ and *T*_2_ = 1*/R*_2_. The spectral bins were reordered in ascending *T*_1_ and *T*_2_ without changing their reconstructed amplitudes. The one-dimensional marginal distributions were obtained by integration (i.e., summation) of the joint distribution over the complementary relaxation-rate dimension:

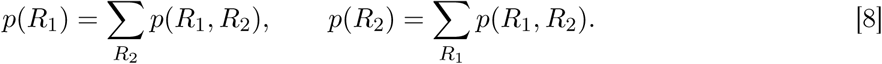

Mean relaxation maps, ⟨*T*_1_⟩ and ⟨*T*_2_⟩, were calculated from the marginal distributions together with the normalized distribution width, defined as the coefficient of variation (CV), *σ_T_*_1_ */*⟨*T*_1_⟩ and *σ_T_*_2_ */*⟨*T*_2_⟩.

Spectral component fractions were calculated by dividing the two-dimensional relaxation space into selected regions. We chose these regions manually and heuristically by inspecting spatial patterns across the complete reconstructed joint relaxation space. Contiguous boundaries on the discrete relaxation grid were then used to group bins with related spatial characteristics. We also examined the resulting component maps for spatial coherence. For each voxel, the component fractions were calculated as

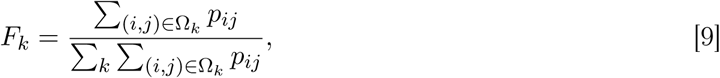

where *p_ij_* denotes the reconstructed probability at grid location (*i, j*), Ω*_k_* denotes the region corresponding to component *k*, and *F_k_* represents its fractional contribution. By construction, Σ*_k_ F_k_* = 1 within each voxel.

We calculated voxel-wise measures of statistical dependence to examine information in the joint *R*_1_-*R*_2_ distribution that is not captured by the individual relaxation dimensions. Covariance and Pearson correlation describe linear dependence between *R*_1_ and *R*_2_. Mutual information (MI) was included as a measure of statistical dependence that does not assume a linear relationship.

These statistical measures were computed directly from the reconstructed joint distribution, *p*(*R*_1_*, R*_2_). Although reconstruction was performed on a logarithmically spaced grid, all statistical moments were calculated using the physical relaxivity values *R*_1_ and *R*_2_. In each voxel, the distribution was normalized such that Σ*_i,j_ p*(*R*_1_*_,i_, R*_2_*_,j_*) = 1.

Covariance was used to quantify the joint linear variation between the two relaxivities:

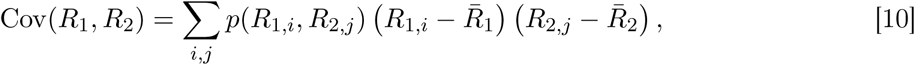

where *R̄*_1_ = Σ*_i,j_ p*(*R*_1*,i*_*, R*_2*,j*_)*R*_1*,i*_ and *R̄*_2_ = Σ*_i,j_ p*(*R*_1*,i*_*, R*_2*,j*_)*R*_2*,j*_ are the mean relaxivities used to center the distribution.

Pearson correlation was used to normalize the covariance by the standard deviations of the corresponding marginal relaxivity distributions:

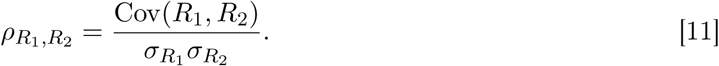

MI was used to quantify statistical dependence between *R*_1_ and *R*_2_ beyond linear correlation. The marginal distributions were defined as *p*(*R*_1_*_,i_*) = Σ*_j_ p*(*R*_1_*_,i_, R*_2_*_,j_*) and *p*(*R*_2_*_,j_*) = Σ*_i_ p*(*R*_1_*_,i_, R*_2_*_,j_*), and MI was calculated as

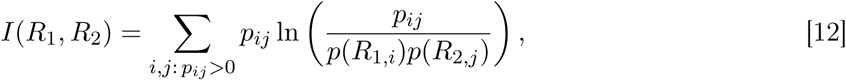

where *p_ij_* = *p*(*R*_1_*_,i_, R*_2_*_,j_*). The natural logarithm was used, so MI is reported in nats. MI is zero when the joint distribution is separable, *p*(*R*_1_*, R*_2_) = *p*(*R*_1_) *p*(*R*_2_), whereas values greater than zero indicate statistical dependence between the two relaxation dimensions.

All statistical measures were calculated from the same normalized, nonnegative *p*(*R*_1_*, R*_2_) distribution within the brain mask. Voxels with zero total spectral probability or invalid numerical values were excluded, and correlation was calculated only when both marginal variances were positive. No *S*_0_ weighting, additional bin-width weighting, or Jacobian transformation was applied because the calculations were performed directly on the native relaxivity reconstruction grid.

## Results

### Phantom Validation

The joint *T*_1_-*T*_2_ fitting approach was evaluated using the UNITY phantom. Figure 2a shows three slices containing the primary quantitative MnCl_2_, PVP, and NiCl_2_ arrays. The joint *T*_1_ maps showed well-defined, spatially uniform compartments across the three phantom slices, with clear variation in relaxation time across the MnCl_2_ and NiCl_2_ concentration series and between the selected PVP compartments. The corresponding joint *T*_2_ maps demonstrated the same overall compartmental organization, although with greater within-compartment spatial variability, particularly in compartments with longer *T*_2_ values.

**Figure 2:**
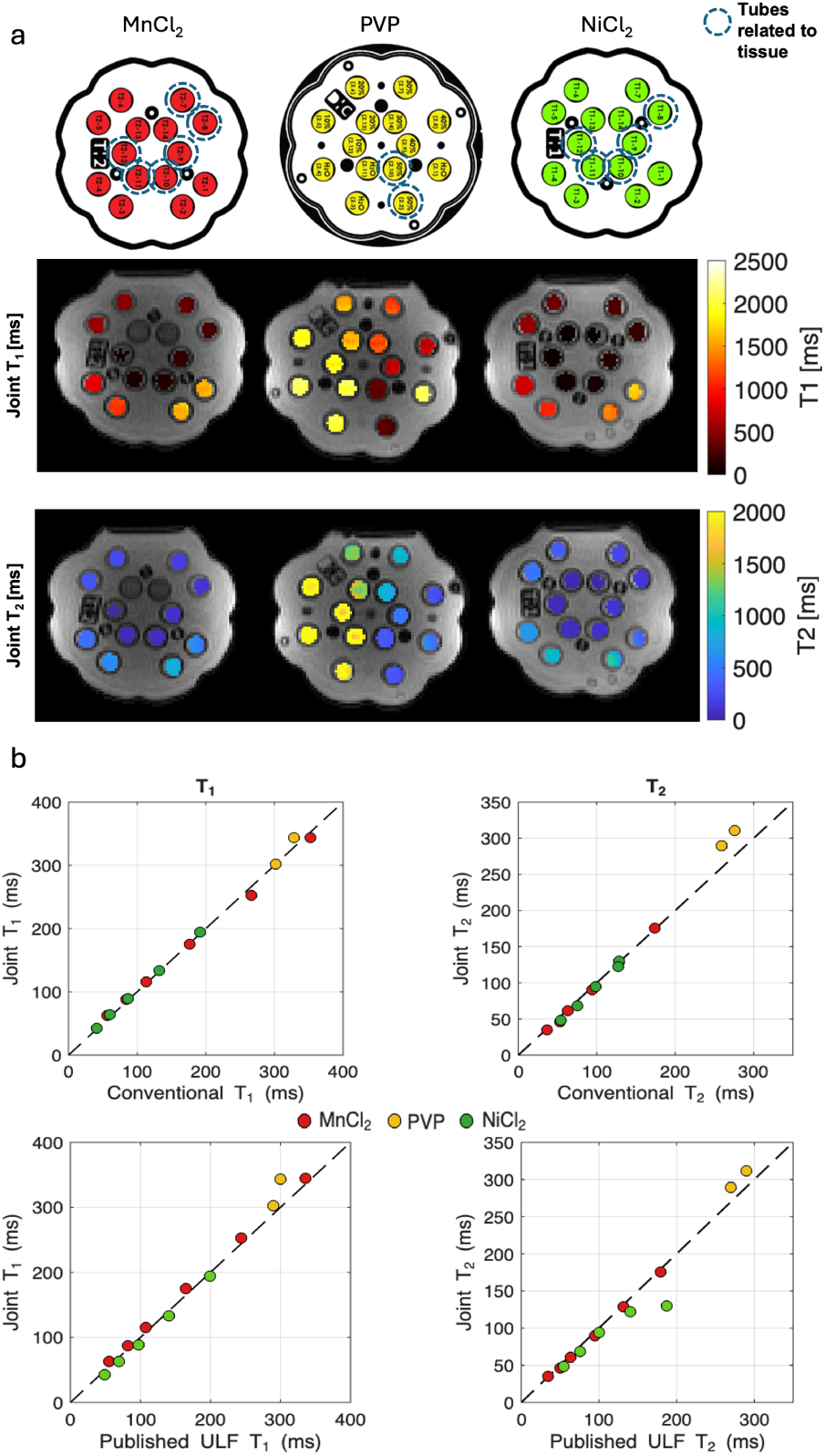
Phantom validation of joint *T*_1_-*T*_2_ relaxation estimates at 0.064 T. (a) Phantom schematics and corresponding joint *T*_1_ and *T*_2_ maps for the three acquired slices containing MnCl_2_, PVP, and NiCl_2_ compartments. Dashed circles indicate compartments with relaxation properties spanning the range relevant to biological tissues and selected for quantitative comparison. (b) Joint *T*_1_ and *T*_2_ estimates compared with mono-exponential relaxometry (top) and previously published MRI measurements of the same phantom acquired at 0.064 T (41) (bottom). Points are colored according to phantom composition: MnCl_2_ (red), PVP (yellow), and NiCl_2_ (green). Dashed lines indicate the line of identity. Compartment-wise mean values, standard errors, and additional published NMR measurements at 0.064 T (42) are provided in the Supplementary Material.

Figure 2b compares the joint relaxation estimates with those obtained using mono-exponential *T*_1_ and *T*_2_ relaxometry. Overall, the two approaches agreed closely across the selected compartments. For example, in the 0.1353 mM MnCl_2_ compartment, mono-exponential fitting yielded *T*_1_ = 352.7 ± 6.2 ms and *T*_2_ = 173.9±1.1 ms, compared with joint-distribution estimates of *T*_1_ = 344.0±5.3 ms and *T*_2_ = 176.0 ± 0.7 ms. Similarly, for the 50% PVP outer compartment, the corresponding estimates were *T*_1_ = 302.2 ±6.1 ms and 302.3 ±6.1 ms, respectively, while the *T*_2_ estimates were 259.3 ±6.0 ms and 289.6 ± 6.1 ms. In the 11.3 mM NiCl_2_ compartment, mono-exponential estimates of *T*_1_ = 131.9 ± 0.5 ms and *T*_2_ = 127.0 ± 0.8 ms were compared with joint estimates of *T*_1_ = 133.4 ± 0.5 ms and *T*_2_ = 122.5 ± 0.7 ms. Joint estimates were also compared with previously published MRI measurements of the same phantom at 0.064 T (41); agreement was generally stronger for *T*_1_, while larger differences were observed for some *T*_2_ compartments. Complete compartment-wise measurements are provided in Supplementary Table S1.

### *In Vivo* Multidimensional Relaxation Imaging

Representative relaxation-weighted images from the multidimensional acquisition are shown in Figure 3. They illustrate both the whole-brain coverage and the range of image contrast obtained with the acquisition. Consecutive axial slices acquired at a single *TI*-*TE* combination show coverage across the imaging volume (Figure 3a), with the highlighted slice selected for subsequent visualization.

**Figure 3:**
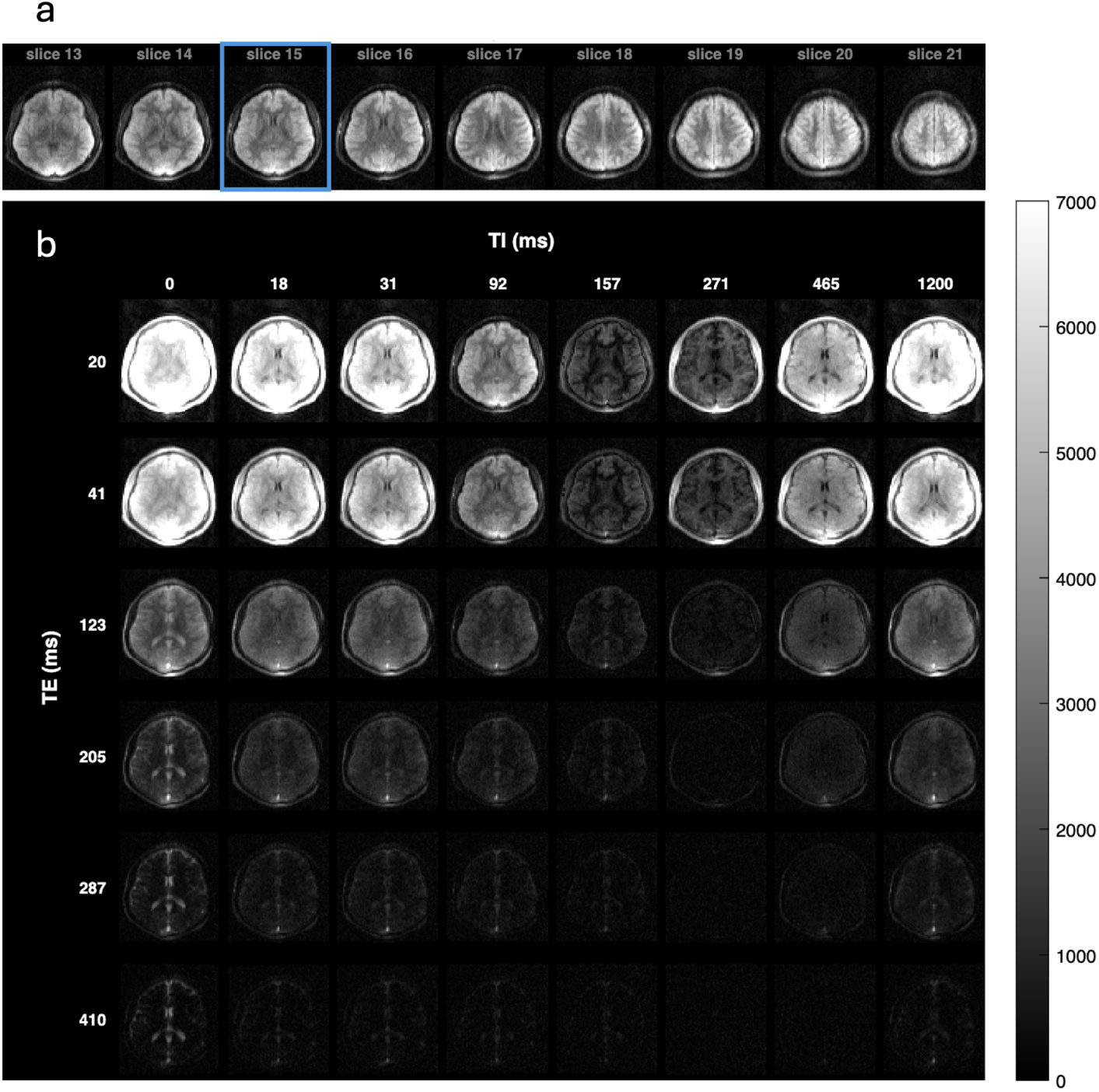
Representative *in vivo* multidimensional relaxation-weighted images acquired using the IR-FSE sequence. (a) Consecutive axial slices demonstrating whole-brain coverage for a representative *TI*-*TE* encoding. The slice outlined in blue was selected for subsequent visualization. (b) Multiple relaxation-weighted images from the selected slice are acquired over representative combinations of inversion time (*TI*) and echo time (*TE*), demonstrating the range of image contrast typically generated by the multidimensional acquisition. Images are displayed using a common grayscale intensity scale.

For this slice, varying *TI* and *TE* produced a broad range of relaxation-weighted signals (Figure 3b), from high signal to progressively attenuated signal approaching the noise level. A range of intermediate tissue contrasts is also visible. Together, these images show that the acquisition samples relaxation weightings across the useful signal range of the multidimensional encoding space.

Mono-exponential voxel-wise *T*_1_ and *T*_2_ maps are shown in Figure 4. These maps show the spatial variation in relaxation times across the brain and serve as a scalar reference for the multidimensional relaxation distributions presented below.

**Figure 4:**
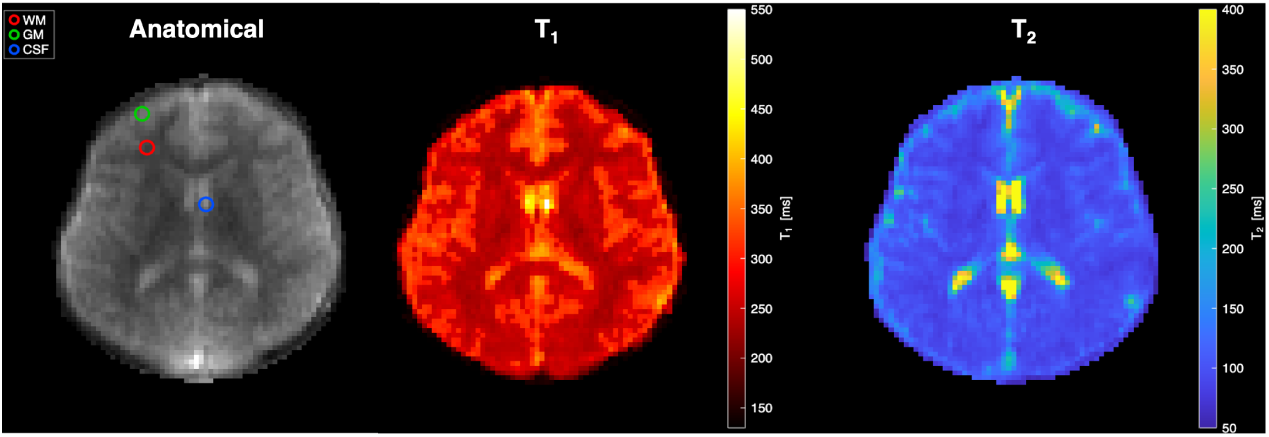
Mono-exponential quantitative relaxation mapping. A representative relaxation-weighted anatomical image is shown with the white matter (WM) (red), gray matter (GM) (green), and cerebrospinal fluid (CSF) (blue) voxels selected for subsequent evaluation, together with voxel-wise mono-exponential *T*_1_ and *T*_2_ maps.

Representative white matter (WM) and gray matter (GM) voxels were selected at locations comparable to those used for low-field relaxation measurements by Jordanova et al. (13). Mono-exponential *T*_1_ estimates were 240 ms in WM and 346 ms in GM, while the corresponding *T*_2_ estimates were 83 ms and 156 ms, respectively. These values were broadly consistent with previously reported relaxation measurements acquired on similar 0.064 T MRI scanners (13; 17).

Figure 5a shows the joint *T*_1_-*T*_2_ relaxation distributions and corresponding marginals for the white matter (WM), gray matter (GM), and cerebrospinal fluid (CSF) voxels identified in Figure 4. The WM and GM spectra were concentrated at shorter and intermediate relaxation times, respectively, while also extending toward longer relaxation components. The CSF distribution was considerably broader and extended toward substantially longer *T*_1_ and *T*_2_ values. Projection of the joint distribution along each relaxation dimension yielded the corresponding *T*_1_ and *T*_2_ marginal distributions.

**Figure 5:**
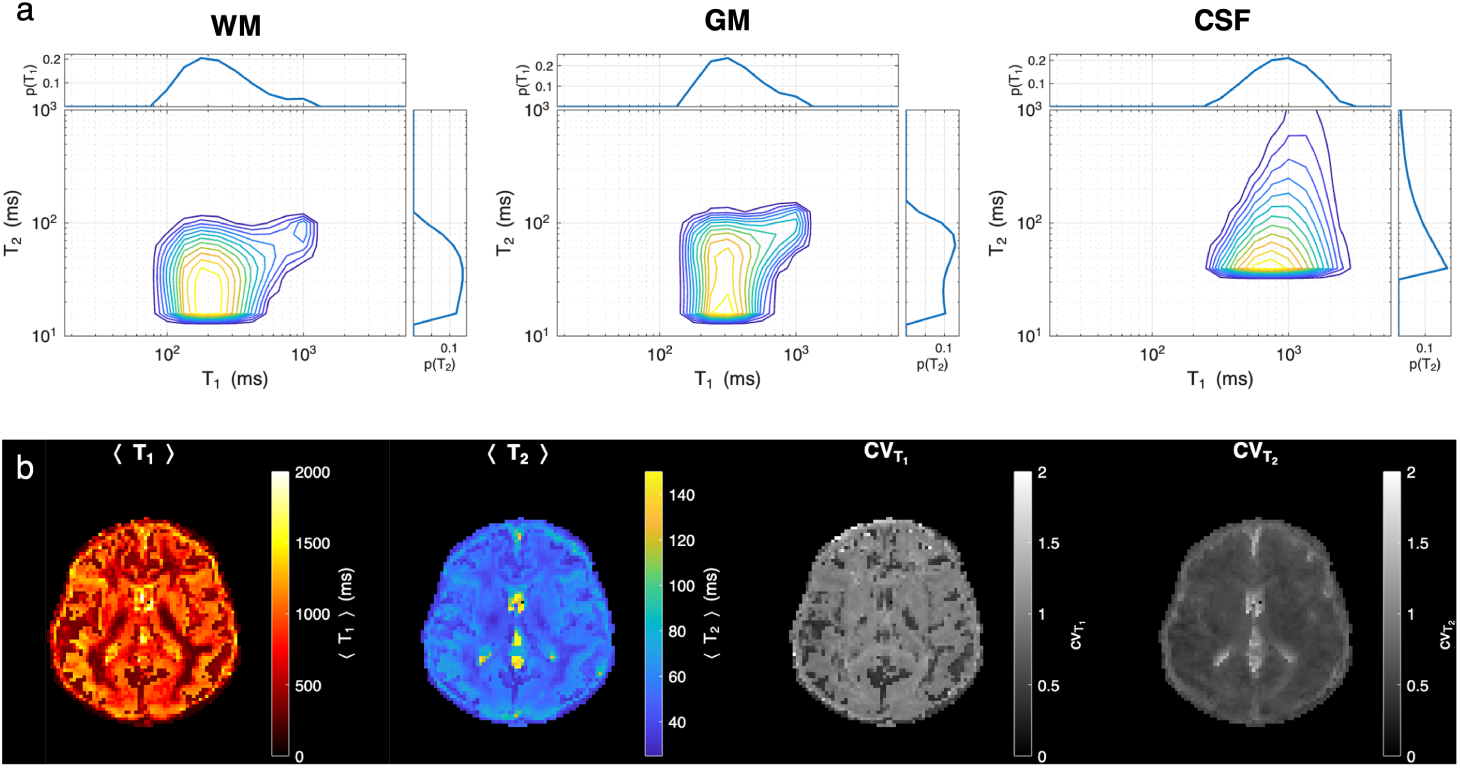
Joint relaxation distributions and distribution-derived summary statistics. (a) Representative joint *T*_1_-*T*_2_ relaxation distributions for white matter (WM), gray matter (GM), and cerebrospinal fluid (CSF) voxels identified in Figure 4. The corresponding *T*_1_ and *T*_2_ marginal distributions, obtained by projection of the joint distribution along the complementary relaxation dimension, are shown above and to the right of each joint distribution, respectively. (b) Voxel-wise mean relaxation times, ⟨*T*_1_⟩ and ⟨*T*_2_⟩, and coefficients of variation, *CV_T_*_1_ and *CV_T_*_2_ , derived from the corresponding marginal distributions. The WM and GM voxels appear to have partial volume (CSF) contamination, which is expected at these voxel sizes.

For the selected voxels, distribution-derived relaxation times are reported as ⟨*T* ⟩ ± *σ_T_* , where *σ_T_* describes the width of the corresponding marginal distribution. The ⟨*T*_1_⟩ ± *σ_T_*_1_ values were 287 ± 198 ms in WM, 396 ± 206 ms in GM, and 1947 ± 626 ms in CSF; the corresponding ⟨*T*_2_⟩ ± *σ_T_*_2_ values were 51 ± 33 ms, 86 ± 61 ms, and 412 ± 281 ms, respectively.

The marginal distributions were further characterized by their mean relaxation time and coefficient of variation, as shown in Figure 5b. The ⟨*T*_1_⟩ and ⟨*T*_2_⟩ maps retain the spatial relaxation contrast while accounting for the full marginal distribution. The coefficient-of-variation maps show how the relative width of these distributions varies across the brain.

Figure 6a shows the complete 21 × 21 *p*(*R*_1_*, R*_2_) reconstruction displayed spatially across the corresponding *T*_1_-*T*_2_ relaxation space. Each location in the grid therefore represents the spatial distribution of the reconstructed spectral probability associated with a particular combination of *T*_1_ and *T*_2_. Summation of the joint distribution along the complementary relaxation dimension yields the corresponding marginal distributions, shown as spatial *T*_1_ marginal maps across the top of the grid and *T*_2_ marginal maps along the left side. The full display can therefore be viewed alongside the marginals to see what is retained, and what is lost, when the joint distribution is projected onto either relaxation dimension alone.

Neighboring spectral bins frequently exhibit similar anatomical contrast, forming contiguous regions of related spatial patterns that vary gradually across the relaxation space and partially overlap with adjacent regions. These patterns are most apparent at intermediate relaxation times. Much less spectral signal is seen near the shortest and longest relaxation times covered by the reconstruction. The apparent inversion efficiency, *η*, which was estimated voxel-wise as part of the multidimensional reconstruction, was close to unity throughout the brain and across the representative slices shown in Figure 6b.

**Figure 6:**
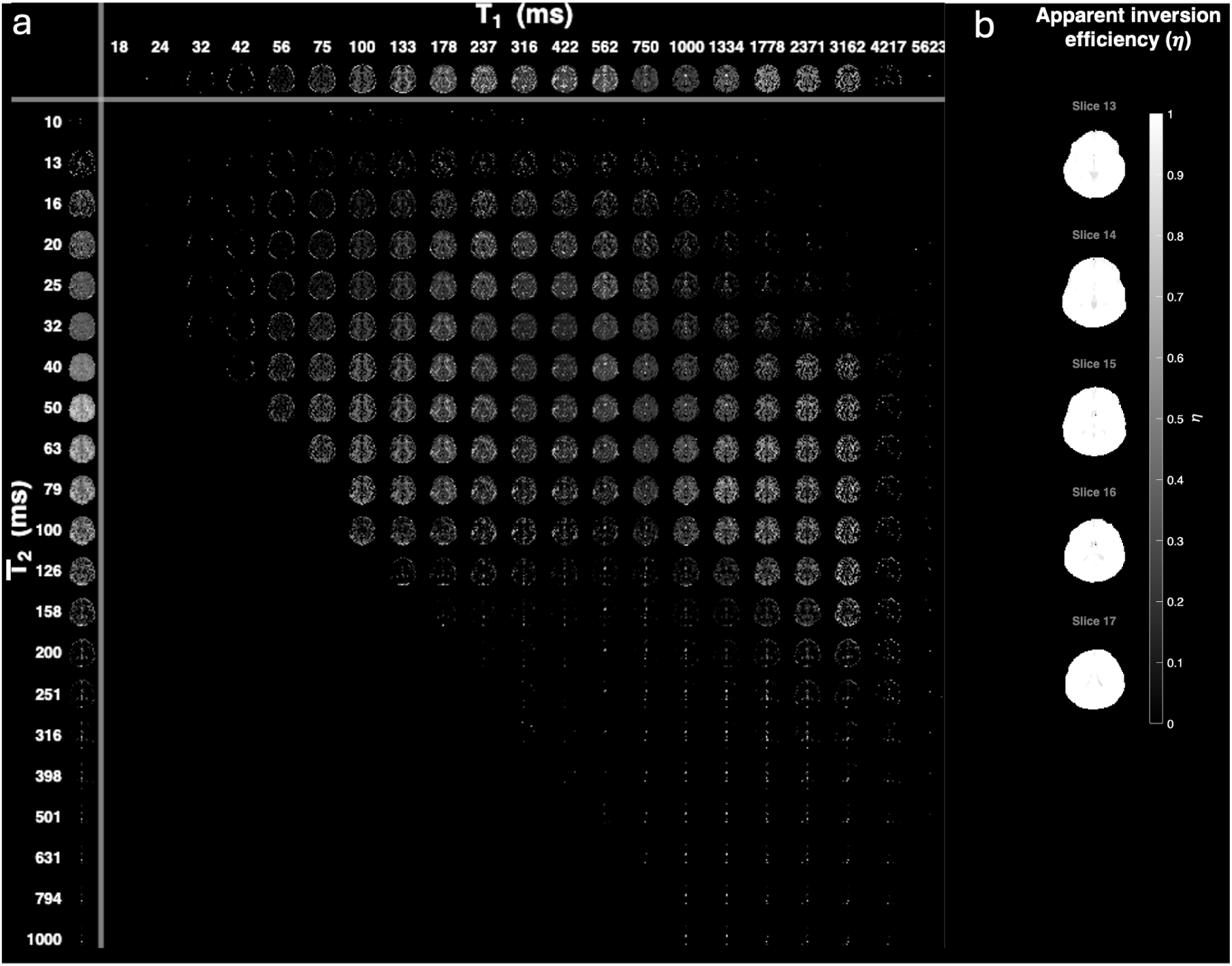
Full spatial representation of the reconstructed joint relaxation distribution and apparent inversion efficiency. (a) Voxel-wise spatial maps of the reconstructed *p*(*T*_1_*, T*_2_) distribution displayed over the complete 21×21 relaxation grid. Each map represents the reconstructed spectral probability associated with the indicated combination of *T*_1_ and *T*_2_. The spatial *T*_1_ marginal distributions, obtained by summation over *T*_2_, are shown across the top, and the corresponding *T*_2_ marginal distributions, obtained by summation over *T*_1_, are shown along the left side. (b) Voxel-wise apparent inversion efficiency (*η*) across five representative axial slices, displayed on a common scale from 0 to 1.

To explore this spatial organization further, Figure 7a shows a compact representation of selected *T*_1_-*T*_2_ bins spanning the relaxation space. Five regions were defined heuristically based on the spatial patterns observed across the full distribution and are indicated by the colored boundaries in the compact representation. For each voxel, the reconstructed probability within the bins assigned to each region was summed and normalized across the five selected regions, producing the component fraction maps shown in Figure 7b. The resulting component maps show distinct spatial patterns across the brain, with different regions of the joint relaxation spectrum contributing different spatial information.

**Figure 7:**
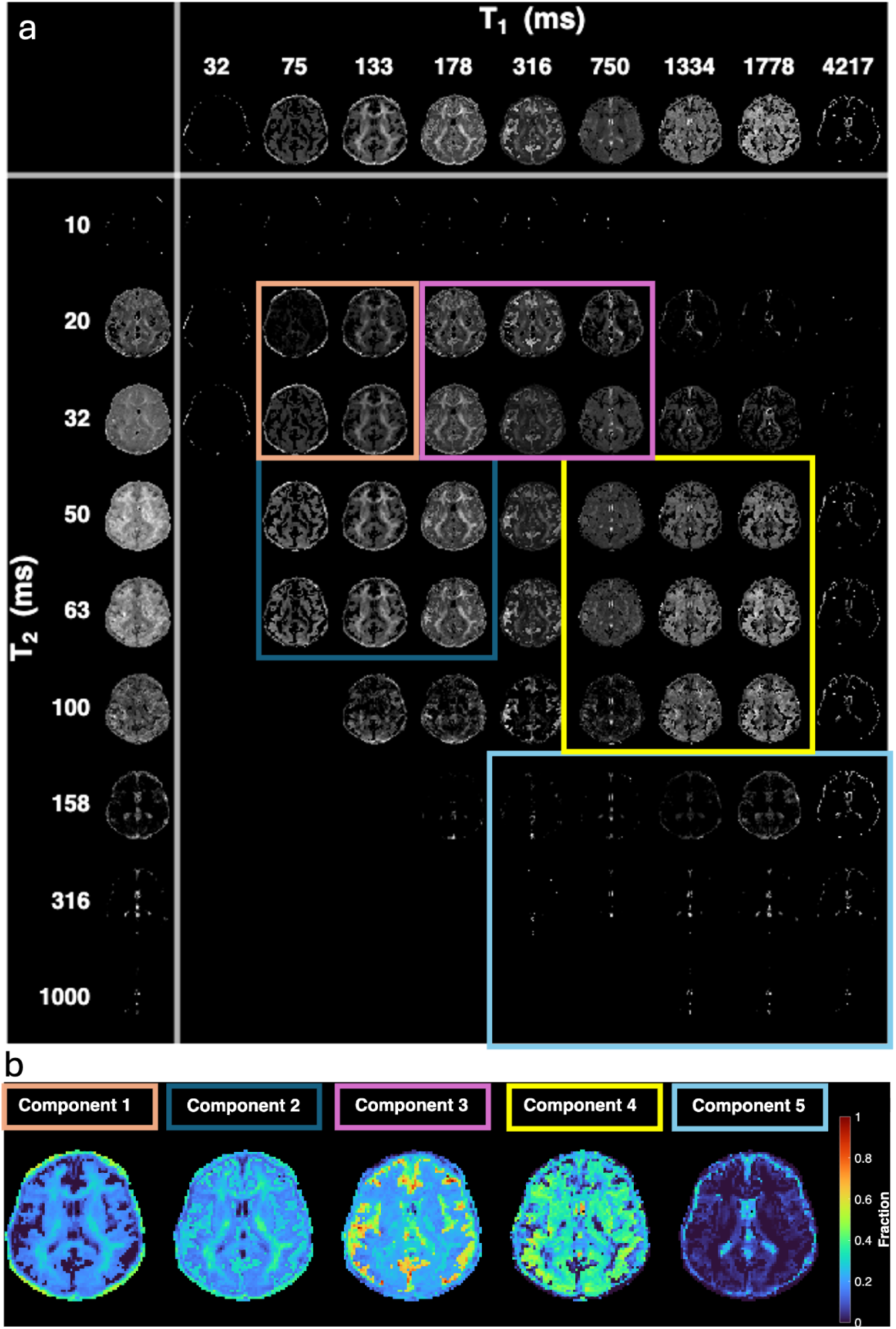
Illustrative decomposition of selected regions within the joint *T*_1_-*T*_2_ relaxation space. (a) Compact representation of selected spectral bins spanning the reconstructed relaxation space. Colored boundaries indicate five regions defined heuristically based on spatial patterns observed across the complete joint distribution shown in Figure 6. (b) Corresponding component fraction maps obtained by summing the reconstructed probability within the bins assigned to each region and normalizing across the five selected regions. Colors identifying the five regions in (a) correspond to the component labels in (b).

### Statistical Characterization

Voxel-wise mutual information (MI), covariance, and Pearson correlation coefficient maps derived from the joint *p*(*R*_1_*, R*_2_) distributions are shown in Figure 8. The three measures exhibited distinct spatial variation across the brain.

**Figure 8:**
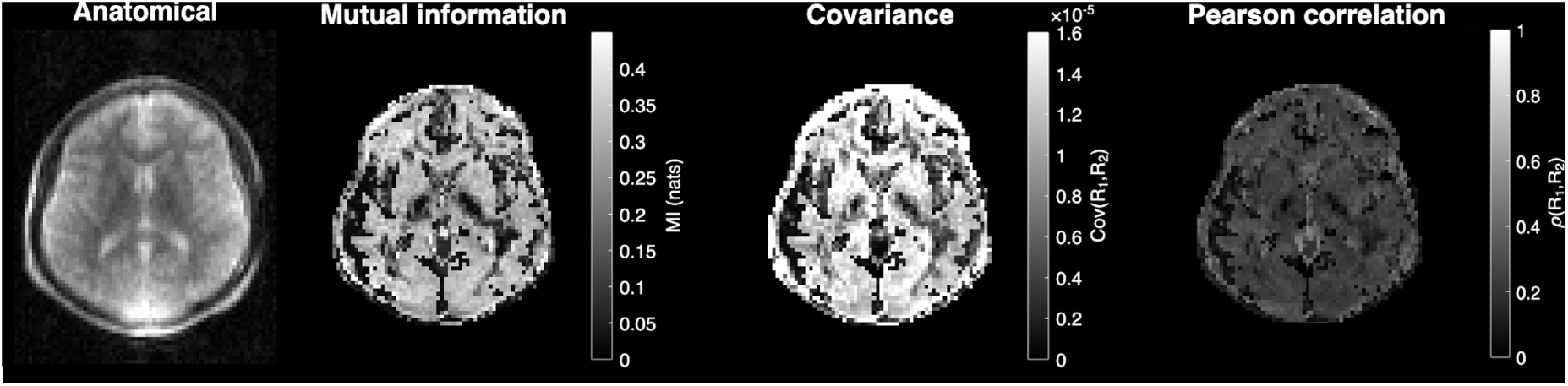
Voxel-wise statistical characterization of the reconstructed joint *p*(*R*_1_*, R*_2_) distributions. A representative relaxation-weighted anatomical image is shown for spatial reference, followed by maps of mutual information (MI), covariance, and Pearson correlation between *R*_1_ and *R*_2_. MI is reported in nats and quantifies statistical dependence between the two relaxation dimensions, while covariance and Pearson correlation characterize linear dependence. Statistical measures were calculated from the normalized, nonnegative joint and marginal probability distributions within the brain mask. The Pearson correlation map is displayed over the range 0-1 for visualization, correlation values were not constrained to be positive.

MI was positive in all valid voxels, with a median value of 0.297, indicating statistical dependence between *R*_1_ and *R*_2_ throughout the brain volume. The median covariance was 1.29 × 10^−5^, while the median Pearson correlation coefficient was 0.254, with positive correlations observed in 98.4% of admissible voxels. The three measures therefore capture different aspects of the relationship between *R*_1_ and *R*_2_, including dependence that is not limited to a linear association.

## Discussion

This study demonstrates the feasibility of whole-brain multidimensional *T*_1_-*T*_2_ relaxation imaging on a portable 0.064 T MRI scanner. We combined a multidimensional IR-FSE acquisition with MADCO reconstruction to obtain joint relaxation distributions *in vivo* within practically achievable scan times. The reconstructed distributions were examined alongside mono-exponential relaxation measurements, distribution-derived metrics, and measures of statistical dependence within the joint relaxation space. To our knowledge, this is the first *in vivo* demonstration of multidimensional *T*_1_-*T*_2_ distribution imaging on a portable ULF MRI scanner. This work is intended as a proof of concept and a starting point for studies addressing more specific biological or clinical questions.

The phantom experiments provide quantitative validation of the multidimensional acquisition and reconstruction framework across a broad range of relaxation properties (Figure 2). Joint *T*_1_ estimates agreed well with both mono-exponential fitting and previously reported measurements at 0.064 T. The *T*_2_ estimates were more variable, particularly in compartments with longer *T*_2_, which may reflect limitations of the multi-echo FSE acquisition. The phantom protocol used LPI = 1 and therefore allowed sampling at short echo times, but the finite echo-train duration limited the longest achievable *TE* and the characterization of slowly decaying components. Repeated refocusing pulses can also generate stimulated-echo pathways and affect the measured signal evolution when refocusing is imperfect (49; 50). Limited control over RF phase cycling and residual odd-even echo variation on the scanner may have further contributed to the variability in *T*_2_. Even with these limitations, the joint estimates were generally consistent with the mono-exponential and previously reported relaxation measurements, supporting the use of the approach for *in vivo* experiments.

The reconstructed relaxation distributions contain information that is not captured by mono-exponential relaxation mapping. The WM and GM mono-exponential estimates (Figure 4) were broadly consistent with previous measurements at 0.064 T (13; 17) and provide a useful reference for interpreting the distributions. However, a mono-exponential fit assigns a single relaxation value to each voxel. The reconstructed spectra show a much broader range of relaxation components at the same locations (Figure 5a). WM and GM distributions were centered at shorter and intermediate relaxation times but often extended toward longer components, while the CSF distribution was particularly broad and shifted toward longer *T*_1_ and *T*_2_. Some of the broader or long-relaxation contributions in nominal WM and GM voxels are likely related to partial-volume effects, given the relatively large voxel dimensions and 5.8 mm slice thickness of the *in vivo* acquisition. CSF is also difficult to characterize with mono-exponential fitting because the maximum *TI* of 1200 ms provides limited direct encoding of its slowly recovering longitudinal magnetization. The multi-dimensional reconstruction instead combines the acquired *TI*, *TE*, and *TR* weightings with the MADCO constraints to estimate probability over a broader joint relaxation domain. It is encouraging that long-relaxation contributions are visible in CSF, but the present data cannot establish their quantitative accuracy outside the most strongly encoded relaxation range.

The full joint relaxation distribution shows clear spatial organization across the two-dimensional relaxation space (Figure 6). The marginal distributions retain much of the relaxation-dependent spatial contrast, but they necessarily collapse one dimension of the joint distribution. This becomes apparent when looking across the full *T*_1_-*T*_2_ space. Neighboring spectral bins often have related spatial patterns and form contiguous regions that change gradually across the distribution. At the same time, bins that contribute to similar locations in one marginal can look quite different depending on their position along the other relaxation dimension. This information cannot be recovered by looking at the marginal distributions alone. The full two-dimensional distribution therefore gives a more complete picture of how the relaxation components are organized and related.

The comparatively low spectral signal toward the shortest and longest relaxation-time limits reflects the relaxation range most strongly encoded by the present acquisition. The sensitivity of the multidimensional measurement to different regions of the relaxation spectrum is therefore expected to depend on the acquisition parameters used to sample the joint space. Future implementations could instead choose the *TI*, *TE*, *TR*, and echo-train sampling based on the region of the relaxation spectrum that is most relevant to a particular application. This would shift sensitivity toward selected relaxation components, rather than trying to sample the entire relaxation space equally.

The component analysis qualitatively explores this spatial organization by grouping selected regions of the joint relaxation space (Figure 7). In the compact display, bins with similar values along one relaxation dimension can have quite different spatial patterns depending on their position along the other dimension (Figure 7a). Grouping some of these regions produces component fraction maps with distinct spatial patterns (Figure 7b). The five components shown here should not be interpreted as definitive tissue compartments or as a systematic segmentation of the relaxation spectrum. They are an illustrative, heuristic decomposition of the information present in the joint space. Some components show patterns that are consistent with known anatomical or relaxation characteristics, including contributions at shorter relaxation times and a long-relaxation component with prominent signal in CSF-containing regions, but we do not assign specific biological identities to them here. Intermediate components may also include contributions from several tissue environments and partial-volume effects, particularly at the spatial resolution of the current ULF acquisition. Future studies will define and validate spectral regions for specific questions, including myelin-sensitive components, tissue transitions, aging-related changes, or pathological alterations, rather than using the illustrative regions selected here.

The statistical analysis gives another way to look at the relationship between the two relaxation dimensions (Figure 8). Mutual information (MI) was positive throughout the analyzed brain, indicating that *R*_1_ and *R*_2_ were not statistically independent. This matters because, if the two dimensions were independent, the joint distribution could be represented by the product of its marginals and there would be little additional information in reconstructing the full distribution, obviating the extra scans required to estimate it. The positive MI values therefore support what is also visible in the spatial maps: the joint distribution contains information that is lost when the two relaxation dimensions are considered separately. MI may also be useful for assessing joint encoding in other multidimensional MRI approaches, including the measurement of distributions that combine relaxation and diffusion dimensions (21; 51).

The observed dependence between the two relaxation dimensions may also have biological relevance. *T*_1_ and *T*_2_ arise from different relaxation mechanisms, but both are influenced by properties of the local tissue environment, including water mobility, macromolecular content, lipid composition, and chemical exchange. Consequently, changes in tissue composition or microstructure may alter not only the individual relaxation distributions but also their relationship within the joint *T*_1_-*T*_2_ space. The statistical measures shown in Figure 8 may therefore be useful for studying how these relationships change with tissue composition, microstructure, or pathology.

## Limitations

A primary limitation of the present implementation is the acquisition time, with the *in vivo* protocol requiring approximately 100 minutes. Further acceleration will be needed for routine clinical imaging. The range of relaxation times directly encoded by the acquisition is also limited. In the phantom, the finite echo-train duration restricted sampling at long *TE*, while the maximum *TI* of 1200 ms in the *in vivo* acquisition provided limited direct encoding of long-*T*_1_ components such as CSF. Components outside the most strongly encoded relaxation range should therefore be interpreted cautiously. Spatial resolution also contributes to partial-volume effects; the relatively large voxel dimensions, including a 5.8 mm slice thickness, likely contribute to the broad distributions and overlapping spectral components seen in Figures 5 and 7. Finally, the current scanner platform provides limited user-level control over several acquisition and reconstruction parameters, including RF and gradient pulse specifications, phase cycling, and aspects of noise correction and image reconstruction. Greater control over these parameters would allow more systematic optimization of multidimensional encoding and effects such as stimulated-echo pathways and residual odd-even echo variation.

The scope of the present study is also limited. The *in vivo* experiment was performed in a single healthy subject and was not designed to assess intersubject variability or reproducibility. Larger studies using standardized acquisition protocols will be needed to address these questions. The spectral component analysis was exploratory, with regions selected manually from spatial and spectral patterns in the reconstructed distributions (Figure 7). These regions were not independently validated as specific tissue or microstructural compartments. More systematic definition and validation of spectral components will therefore be needed before they can be interpreted as quantitative biological markers.

## Future Directions

Future work should focus on improving both the acquisition efficiency and the biological specificity of multidimensional relaxation imaging at ULF. Shorter acquisition times would improve practical feasibility, while the relaxation encoding could be broadened or targeted toward specific regions of the spectrum depending on the application. Further acceleration and development of denoising strategies, including approaches that make use of both magnitude and phase information (52), may also increase the efficiency and robustness of multidimensional imaging at ULF. Greater control over RF and gradient waveforms, potentially through open source sequence development frameworks such as Pulseq (53), would also enable further optimization of the multidimensional acquisition.

A second priority is the systematic identification and validation of biologically meaningful spectral features. The short-*T*_1_ and short-*T*_2_ regions observed within the joint relaxation space may be particularly relevant to future investigation of myelin-sensitive components, while other spectral regions could be defined to address specific biological or clinical questions. Larger studies will also be needed to characterize normative multidimensional relaxation distributions and their variability across populations. The framework could also be extended to additional encoding dimensions, particularly diffusion. This could enable higher-dimensional distributions such as *p*(*T*_1_*, T*_2_*, MD*) (21) or related joint relaxation-diffusion representations (51), with the potential to increase the specificity of multidimensional MRI at ULF.

## Broader Implications

Extending quantitative and multidimensional MRI methods to portable ULF scanners could broaden the information available from accessible neuroimaging beyond anatomical contrast alone. The present whole-brain joint *T*_1_-*T*_2_ demonstration is an initial step toward bringing multidimensional tissue characterization to lower-cost, portable ULF MRI scanners.

## Conclusions

This work demonstrates the feasibility of whole-brain multidimensional *T*_1_-*T*_2_ relaxation imaging on a portable 0.064 T MRI scanner. Joint relaxation distributions were reconstructed using an IR-FSE acquisition and MADCO, with quantitative phantom measurements used for validation before applying the approach *in vivo*. The joint distributions contained information beyond the single relaxation values obtained from mono-exponential mapping. This included the marginal distributions and their derived measures, spatial patterns across the joint relaxation space, and measurable dependence between the two relaxation dimensions. To our knowledge, this is the first *in vivo* demonstration of multidimensional *T*_1_-*T*_2_ distribution imaging on a portable ULF MRI scanner. Our results support further development of multidimensional MRI for biological and clinical applications.

## Acknowledgments

This study was supported by the Intramural Research Program (IRP) of the *Eunice Kennedy Shriver* National Institute of Child Health and Human Development under award 1-ZIA-HD008971-07, the National Institute of Neurological Diseases and Stroke Intramural Research Program under award ZIC-NS009450, and the National Institute on Aging. This work was also funded by the Department of War in the Military Traumatic Brain Injury Initiative (MTBI^2^) under award, HU0001-22-2-0058. This work utilized computational resources of the NIH HPC Biowulf cluster (http://hpc.nih.gov). The authors have no conflicts of interest to declare. The views, information or content, and conclusions presented do not necessarily represent the official position or policy of, nor should any official endorsement be inferred on the part of, the Uniformed Services University, the Department of War, the U.S. Government, or the Henry M. Jackson Foundation for the Advancement of Military Medicine, Inc. The contributions of the NIH author(s) are considered Works of the United States Government. The findings and conclusions presented in this paper are those of the author(s) and do not necessarily reflect the views of the NIH or the U.S. Department of Health and Human Services. The authors wish to thank Sean Deoni of the Gates Foundation and the UNITY consortium created under the auspices of the Gates Foundation for their continued leadership and support in advancing ULF MRI developments and applications in LMICs.

## References

1 Cooley CZ, McDaniel PC, Stockmann JP, Srinivas SA, Cauley SF, Śliwiak M, Sappo CR, Vaughn CF, Guerin B, Rosen MS, Lev MH, Wald LL. 2021. A portable scanner for magnetic resonance imaging of the brain. Nature Biomedical Engineering 5:229–239. doi: 10.1038/s41551-020-00641-5.

2 Mazurek MH, Cahn BA, Yuen MM, Prabhat AM, Chavva IR, Shah JT, Crawford AL, Welch EB, Rothberg J, Sacolick L, Poole M, Wira C, Matouk CC, Ward A, Timario N, Leasure A, Beekman R, Peng TJ, Witsch J, Antonios JP, Falcone GJ, Gobeske KT, Petersen N, Schindler J, Sansing L, Gilmore EJ, Hwang DY, Kim JA, Malhotra A, Sze G, Rosen MS, Kimberly WT, Sheth KN. 2021. Portable, bedside, low-field magnetic resonance imaging for evaluation of intracerebral hemorrhage. Nature Communications 2021 12:1 12:5119–. doi:10.1038/s41467-021-25441-6.

3 Basser P. 2022. Detection of stroke by portable, low-field mri: A milestone in medical imaging. Science Advances 8:eabp9307. doi:10.1126/sciadv.abp9307.

4 Balaji S, Wiley N, Poorman ME, Kolind SH. 2024. Low-field mri for use in neurological diseases. Current Opinion in Neurology 37:381–391. doi:10.1097/wco.0000000000001282.

5 Yuen MM, Prabhat AM, Mazurek MH, Chavva IR, Crawford A, Cahn BA, Beekman R, Kim JA, Gobeske KT, Petersen NH, Falcone GJ, Gilmore EJ, Hwang DY, Jasne AS, Amin H, Sharma R, Matouk C, Ward A, Schindler J, Sansing L, de Havenon A, Aydin A, Wira C, Sze G, Rosen MS, Kimberly WT, Sheth KN. 2022. Portable, low-field magnetic resonance imaging enables highly accessible and dynamic bedside evaluation of ischemic stroke. Science Advances 8. doi: 10.1126/sciadv.abm3952.

6 Chetcuti K, Chilingulo C, Goyal MS, Vidal L, O’Brien NF, Postels DG, Seydel KB, Taylor TE. 2022. Implementation of a low-field portable mri scanner in a resource-constrained environment: Our experience in malawi. American Journal of Neuroradiology 43:670–674. doi:10.3174/ajnr.A7494.

7 Arnold TC, Tu D, Okar SV, Nair G, By S, Kawatra KD, Robert-Fitzgerald TE, Desiderio LM, Schindler MK, Shinohara RT, Reich DS, Stein JM. 2022. Sensitivity of portable low-field magnetic resonance imaging for multiple sclerosis lesions. NeuroImage: Clinical 35:103101. doi: 10.1016/j.nicl.2022.103101.

8 Kazimuddin HF, Pathakamuri A, Yi J, Soldatos A, Turtzo LC, Latour LL, Chittiboina P, Brown DA, Reich DS, Horovitz SG. 2026. Gadolinium-enhanced portable ultra-low-field mri for evaluating various intracranial pathologies. American Journal of Neuroradiology 47:1927–1932. doi: 10.3174/ajnr.A9249.

9 Abate F, Adu-Amankwah A, Ae-Ngibise KA, Agbokey F, Agyemang VA, Agyemang CT, Akgun C, Ametepe J, Arichi T, Asante KP, Balaji S, Baljer L, Basser PJ, Beauchemin J, Bennallick C, Berhane Y, Boateng-Mensah Y, Bourke NJ, Bradford L, Bruchhage MM, Lorente RC, Cawley P, Cercignani M, Sa VD, de Canha A, de Navarro N, D C ID, Delarosa J, Donald KA, Dvorak A, Edwards AD, Field D, Frail H, Freeman B, George T, Gholam J, Guerrero-Gonzalez J, Hajnal JV, Haque R, Hollander W, Hoodbhoy Z, Huentelman M, Jafri SK, Jones DK, Joubert F, Karaulanov T, Kasaro MP, Knackstedt S, Kolind S, Koshy B, Kravitz R, Lafayette SL, Lee AC, Lena B, Lepore N, Linguraru M, Ljungberg E, Lockart Z, Loth E, Mannam P, Masemola KM, Moran R, Murphy D, Nakwa FL, Nankabirwa V, Nelson CA, North K, Nyame S, Halloran RO, O’Muircheartaigh J, Oakley BF, Odendaal H, Ongeti CM, Onyango D, Oppong SA, Padormo F, Parvez D, Paus T, Pepper MS, Phiri KS, Poorman M, Ringshaw JE, Rogers J, Rutherford M, Sabir H, Sacolick L, Seal M, Sekoli ML, Shama T, Siddiqui K, Sindano N, Spelke MB, Springer PE, Suleman FE, Sundgren PC, Teixeira R, Terekegn W, Traughber M, Tuuli MG, van Rensburg J, Váša F, Velaphi S, Velasco P, Viljoen IM, Vokhiwa M, Webb A, Weiant C, Wiley N, Wintermark P, Yibetal K, Deoni SC, Williams SC. 2024. Unity: A low-field magnetic resonance neuroimaging initiative to characterize neurodevelopment in low and middle-income settings. Developmental Cognitive Neuroscience 69:101397. doi:10.1016/j.dcn.2024.101397.

10 Ma D, Gulani V, Seiberlich N, Liu K, Sunshine JL, Duerk JL, Griswold MA. 2013. Magnetic resonance fingerprinting. Nature 495:187–192. doi:10.1038/nature11971.

11 Filo S, Shtangel O, Salamon N, Kol A, Weisinger B, Shifman S, Mezer AA. 2019. Disentangling molecular alterations from water-content changes in the aging human brain using quantitative mri. Nature Communications 10:3403. doi:10.1038/s41467-019-11319-1.

12 Rooney WD, Johnson G, Li X, Cohen ER, Kim SG, Ugurbil K, Springer CS. 2007. Magnetic field and tissue dependencies of human brain longitudinal 1h2o relaxation in vivo. Magnetic Resonance in Medicine 57:308–318. doi:10.1002/mrm.21122.

13 Jordanova KV, Martin MN, Ogier SE, Poorman ME, Keenan KE. 2023. In vivo quantitative mri: T1 and t2 measurements of the human brain at 0.064 t. Magnetic Resonance Materials in Physics, Biology and Medicine 36:487–498. doi:10.1007/s10334-023-01095-x.

14 O’Reilly T, Webb AG. 2022. In vivo t1 and t2 relaxation time maps of brain tissue, skeletal muscle, and lipid measured in healthy volunteers at 50 mt. Magnetic Resonance in Medicine 87:884–895. doi:10.1002/mrm.29009.

15 Deoni SC, O’Muircheartaigh J, Ljungberg E, Huentelman M, Williams SC. 2022. Simultaneous high-resolution t2-weighted imaging and quantitative t2 mapping at low magnetic field strengths using a multiple te and multi-orientation acquisition approach. Magnetic Resonance in Medicine 88:1273–1281. doi:10.1002/mrm.29273.

16 Padormo F, Cawley P, Dillon L, Hughes E, Almalbis J, Robinson J, Maggioni A, Botella MDLF, Cromb D, Price A, Arlinghaus L, Pitts J, Luo T, Zhang D, Deoni SC, Williams S, Malik S, O’Muircheartaigh J, Counsell SJ, Rutherford M, Arichi T, Edwards AD, Hajnal JV. 2023. In vivo t1 mapping of neonatal brain tissue at 64 mt. Magnetic Resonance in Medicine 89:1016–1025. doi:10.1002/mrm.29509.

17 Lena B, Padormo F, Teixeira RPA, Bennallick C, Gholam J, van den Broek R, Lafayette SL, Vavasour I, Cercignani M, Jones DK, Kolind S, Hajnal J, Bourke N, Dong Y, Hollander WJ, Karaulanov T, Deoni SC, Williams SC, Sundgren PC, Webb AG, Ljungberg E. 2025. Repeatability and reproducibility of rapid t1 mapping of brain tissues at 64 mt: A multicentre study. Imaging Neuroscience 3. doi:10.1162/imag.a.916.

18 Whittall KP, MacKay AL, Graeb DA, Nugent RA, Li DK, Paty DW. 1997. In vivo measurement of t2 distributions and water contents in normal human brain. Magnetic Resonance in Medicine 37:34–43. doi:10.1002/mrm.1910370107.

19 Labadie C, Lee JH, Rooney WD, Jarchow S, Aubert-Frécon M, Springer CSJ, Möller HE. 2014. Myelin water mapping by spatially regularized longitudinal relaxographic imaging at high magnetic fields. Magnetic Resonance in Medicine 71:375–387. doi:10.1002/mrm.24670.

20 Avram AV, Sarlls JE, Basser PJ. 2019. Measuring non-parametric distributions of intravoxel mean diffusivities using a clinical mri scanner. NeuroImage 185:255–262. doi:10.1016/j.neuroimage.2018.10.030.

21 Benjamini D, Bouhrara M, Komlosh ME, Iacono D, Perl DP, Brody DL, Basser PJ. 2021. Multidimensional mri for characterization of subtle axonal injury accelerated using an adaptive nonlocal multispectral filter. Frontiers in Physics 9:737374. doi:10.3389/fphy.2021.737374.

22 Slator PJ, Palombo M, Miller KL, Westin CF, Laun F, Kim D, Haldar JP, Benjamini D, Lember-skiy G, de Almeida Martins JP, Hutter J. 2021. Combined diffusion-relaxometry microstructure imaging: Current status and future prospects. Magnetic Resonance in Medicine 86:2987–3011. doi:10.1002/mrm.28963.

23 English AE, Whittall KP, Joy ML, Henkelman RM. 1991. Quantitative two-dimensional time correlation relaxometry. Magnetic Resonance in Medicine 22:425–434. doi:10.1002/mrm.1910220250.

24 Kim D, Doyle EK, Wisnowski JL, Kim JH, Haldar JP. 2017. Diffusion-relaxation correlation spectroscopic imaging: A multidimensional approach for probing microstructure. Magnetic Resonance in Medicine 78:2236–2249. doi:10.1002/mrm.26629.

25 Barsoum S, Latimer CS, Nolan AL, Barrett A, Chang K, Troncoso JC, Keene CD, Benjamini D. 2025. Multidimensional mri reveals cortical astrogliosis linked to dementia in alzheimer’s disease. Brain Communications 7:fcaf245. doi:10.1093/braincomms/fcaf245.

26 Kundu S, Barsoum S, Ariza J, Nolan AL, Latimer CS, Keene CD, Basser PJ, Benjamini D. 2023. Mapping the individual human cortex using multidimensional mri and unsupervised learning. Brain Communications 5. doi:10.1093/braincomms/fcad258.

27 Park JS, Manninen E, Yang Y, Benjamini D. 2026. Informed dictionary-guided monte carlo inversion for robust and reproducible multidimensional mri. Magnetic Resonance in Medicine 95:2947–2962. doi:10.1002/mrm.70228.

28 Avram AV, Sarlls JE, Basser PJ. 2021. Whole-brain imaging of subvoxel t1-diffusion correlation spectra in human subjects. Frontiers in Neuroscience 15:671465. doi:10.3389/fnins.2021.671465.

29 Magdoom KN, Komlosh ME, Saleem K, Gasbarra D, Basser PJ. 2022. High resolution ex vivo diffusion tensor distribution mri of neural tissue. Frontiers in Physics 10:807000. doi: 10.3389/fphy.2022.807000.

30 Slator PJ, Hutter J, Palombo M, Jackson LH, Ho A, Panagiotaki E, Chappell LC, Rutherford MA, Hajnal JV, Alexander DC. 2019. Combined diffusion-relaxometry mri to identify dysfunction in the human placenta. Magnetic Resonance in Medicine 82:95–106. doi:10.1002/mrm.27733.

31 Benjamini D, Priemer DS, Perl DP, Brody DL, Basser PJ. 2023. Mapping astrogliosis in the individual human brain using multidimensional mri. Brain 146:1212–1226. doi:10.1093/brain/awac298.

32 Benjamini D, Iacono D, Komlosh ME, Perl DP, Brody DL, Basser PJ. 2021. Diffuse axonal injury has a characteristic multidimensional mri signature in the human brain. Brain 144:800–816. doi:10.1093/brain/awaa447.

33 de Almeida Martins JP, Tax CM, Szczepankiewicz F, Jones DK, Westin CF, Topgaard D. 2020. Transferring principles of solid-state and laplace nmr to the field of in vivo brain mri. Magnetic Resonance 1:27–43. doi:10.5194/mr-1-27-2020.

34 Martin J, Reymbaut A, Schmidt M, Doerfler A, Uder M, Laun FB, Topgaard D. 2021. Non-parametric d-r1-r2 distribution mri of the living human brain. NeuroImage 245:118753. doi: 10.1016/j.neuroimage.2021.118753.

35 Johnson JT, Irfanoglu MO, Manninen E, Ross TJ, Yang Y, Laun FB, Martin J, Topgaard D, Benjamini D. 2024. In vivo disentanglement of diffusion frequency-dependence, tensor shape, and relaxation using multidimensional mri. Human Brain Mapping 45:e26697. doi:10.1002/hbm.26697.

36 Bai R, Benjamini D, Cheng J, Basser PJ. 2016. Fast, accurate 2d-mr relaxation exchange spectroscopy (rexsy): Beyond compressed sensing. Journal of Chemical Physics 145:154202. doi:10.1063/1.4964144.

37 Callaghan PT. 2011. Translational dynamics and magnetic resonance: Principles of pulsed gradient spin echo NMR. Oxford University Press.

38 Bai R, Cloninger A, Czaja W, Basser PJ. 2015. Efficient 2d mri relaxometry using compressed sensing. Journal of Magnetic Resonance 255:88–99. doi:10.1016/j.jmr.2015.04.002.

39 Benjamini D, Basser PJ. 2016. Use of marginal distributions constrained optimization (madco) for accelerated 2d mri relaxometry and diffusometry. Journal of Magnetic Resonance 271:40–45. doi:10.1016/j.jmr.2016.08.004.

40 Venkataramanan L, Song YQ, Hürlimann MD. 2002. Solving fredholm integrals of the first kind with tensor product structure in 2 and 2.5 dimensions. IEEE Transactions on Signal Processing 50:1017–1026. doi:10.1109/78.995059.

41 Ljungberg E, Padormo F, Poorman M, Clemensson P, Bourke N, Evans JC, Gholam J, Vavasour I, Kollind SH, Lafayette SL, Bennallick C, Donald KA, Bradford LE, Lena B, Vokhiwa M, Shama T, Siew J, Sekoli L, van Rensburg J, Pepper MS, Khan A, Madhwani A, Banda FA, Mwila ML, Cassidy AR, Moabi K, Sephi D, Boakye RA, Ae-Ngibise KA, Asante KP, Hollander WJ, Karaulanov T, Williams SC, Deoni S. 2025. Characterization of portable ultra-low field mri scanners for multi-center structural neuroimaging. Human Brain Mapping 46:e70217. doi: 10.1002/hbm.70217.

42 Martin MN, Jordanova KV, Kos AB, Russek SE, Keenan KE, Stupic KF. 2023. Relaxation measurements of an mri system phantom at low magnetic field strengths. Magnetic Resonance Materials in Physics, Biology and Medicine 36:477–485. doi:10.1007/s10334-023-01086-y.

43 Veraart J, Novikov DS, Christiaens D, Ades-Aron B, Sijbers J, Fieremans E. 2016. Denoising of diffusion mri using random matrix theory. NeuroImage 142:394–406. doi:10.1016/j.neuroimage.2016.08.016.

44 Garyfallidis E, Brett M, Amirbekian B, Rokem A, van der Walt S, Descoteaux M, Nimmo-Smith I, Contributors D. 2014. Dipy, a library for the analysis of diffusion mri data. Frontiers in Neuroinformatics 8:8. doi:10.3389/fninf.2014.00008.

45 Jenkinson M, Smith S. 2001. A global optimisation method for robust affine registration of brain images. Medical Image Analysis 5:143–156. doi:10.1016/s1361-8415(01)00036-6.

46 Jenkinson M, Bannister P, Brady M, Smith S. 2002. Improved optimization for the robust and accurate linear registration and motion correction of brain images. NeuroImage 17:825–841. doi:10.1006/nimg.2002.1132.

47 Smith SM, Jenkinson M, Woolrich MW, Beckmann CF, Behrens TE, Johansen-Berg H, Bannister PR, Luca MD, Drobnjak I, Flitney DE, Niazy RK, Saunders J, Vickers J, Zhang Y, Stefano ND, Brady JM, Matthews PM. 2004. Advances in functional and structural mr image analysis and implementation as fsl. NeuroImage 23:S208–S219. doi:10.1016/j.neuroimage.2004.07.051.

48 Bakker CJ, de Graaf CN, van Dijk P. 1984. Restoration of signal polarity in a set of inversion recovery nmr images. IEEE Transactions on Medical Imaging 3:197–202. doi:10.1109/tmi.1984.4307681.

49 Song YQ. 2002. Categories of coherence pathways for the cpmg sequence. Journal of Magnetic Resonance 157:82–91. doi:10.1006/jmre.2002.2577.

50 Le Roux P. 2002. Non-CPMG Fast Spin Echo with Full Signal. Journal of Magnetic Resonance 155:278–292. doi:10.1006/JMRE.2002.2523.

51 Magdoom KN, Sarlls JE, Basser PJ. 2026. Towards mesoscopic human brain imaging using non-parametric diffusion tensor distribution (dtd) mri. bioRxiv doi:10.64898/2026.04.12.718025.

52 Bai R, Koay CG, Hutchinson E, Basser PJ. 2014. A framework for accurate determination of the t2 distribution from multiple echo magnitude mri images. Journal of Magnetic Resonance 244:53–63. doi:10.1016/j.jmr.2014.04.016.

53 Layton KJ, Kroboth S, Jia F, Littin S, Yu H, Leupold J, Nielsen JF, Stöcker T, Zaitsev M. 2017. Pulseq: A rapid and hardware-independent pulse sequence prototyping framework. Magnetic Resonance in Medicine 77:1544–1552. doi:10.1002/mrm.26235.

