## Supplemental Table 1 for "Multidimensional *T*_1_-*T*_2_ Relaxation Imaging *In Vivo* on a Portable 0.064 T MRI Scanner"

### Table S1

Quantitative MRI phantom relaxation data summary for individual $T_1$ and $T_2$ fitting and joint $T_1$-$T_2$ fitting. MRI reference values are from previously reported 0.064 T measurements [1], and NMR reference values are shown where available [2].

| Tube | Content | Conc. (mM) | Mono-exponential $T_1$ (ms) Mean | SE | Mono-exponential $T_2$ (ms) Mean | SE | Joint distribution $T_1$ (ms) Mean | SE | Joint distribution $T_2$ (ms) Mean | SE | MRI reference $T_1$ (ms) Mean | MRI reference $T_2$ (ms) Mean | NMR reference $T_1$ (ms) Mean | SE | NMR reference $T_2$ (ms) Mean | SE |
|---|---|---|---|---|---|---|---|---|---|---|---|---|---|---|---|---|
| 7 | $MnCl_2$ | 0.135 | 352.66 | 6.16 | 173.92 | 1.06 | 344.01 | 5.33 | 176.00 | 0.67 | 336 | 180 | 343.27 | 0.53 | 185.37 | 2.04 |
| 8 | $MnCl_2$ | 0.193 | 266.63 | 3.15 | 128.86 | 0.81 | 252.29 | 4.35 | 128.85 | 0.64 | 244 | 131 | — | — | — | — |
| 9 | $MnCl_2$ | 0.277 | 177.03 | 0.92 | 94.42 | 1.22 | 175.64 | 0.97 | 89.95 | 0.79 | 165 | 95 | 168.03 | 0.65 | 88.98 | 0.29 |
| 10 | $MnCl_2$ | 0.428 | 112.71 | 0.43 | 63.51 | 1.11 | 115.52 | 0.44 | 60.93 | 0.96 | 108 | 64 | — | — | — | — |
| 11 | $MnCl_2$ | 0.556 | 84.26 | 0.56 | 52.87 | 0.87 | 87.80 | 0.88 | 46.41 | 0.87 | 82 | 50 | 78.74 | 0.40 | 44.18 | 0.32 |
| 12 | $MnCl_2$ | 0.790 | 56.34 | 0.77 | 36.45 | 1.34 | 62.97 | 0.92 | 34.93 | 0.62 | 56 | 35 | — | — | — | — |
| 2 | PVP (O) | 50% | 302.22 | 6.13 | 259.34 | 5.98 | 302.26 | 6.10 | 289.63 | 6.14 | 290 | 270 | — | — | — | — |
| 10 | PVP (I) | 50% | 328.35 | 4.26 | 276.09 | 3.56 | 343.81 | 3.31 | 311.03 | 3.93 | 300 | 290 | — | — | — | — |
| 8 | $NiCl_2$ | 7.74 | 191.44 | 0.77 | 129.01 | 1.27 | 194.84 | 0.40 | 129.76 | 1.05 | 200 | 187 | — | — | — | — |
| 9 | $NiCl_2$ | 11.3 | 131.94 | 0.54 | 126.99 | 0.76 | 133.38 | 0.51 | 122.47 | 0.70 | 141 | 141 | 138.70 | 0.29 | 137.60 | 0.36 |
| 10 | $NiCl_2$ | 16.5 | 86.66 | 0.55 | 98.69 | 1.03 | 89.14 | 0.80 | 94.23 | 0.83 | 97 | 100 | — | — | — | — |
| 11 | $NiCl_2$ | 23.3 | 60.43 | 0.62 | 75.74 | 1.21 | 63.65 | 0.99 | 68.32 | 0.94 | 69 | 76 | 69.79 | 0.22 | 70.98 | 0.24 |
| 12 | $NiCl_2$ | 32.7 | 41.39 | 0.42 | 54.70 | 0.80 | 43.01 | 0.57 | 48.02 | 0.47 | 49 | 55 | — | — | — | — |

*Notes:* Concentrations are reported in mM except for PVP, which is reported as a percentage. O = outer; I = inner. MRI and NMR columns contain reference relaxation values from the indicated sources. "—" indicates that no reference value was available.

### References

1. Ljungberg E, Padormo F, Poorman M, et al. Characterization of portable ultra-low field MRI scanners for multi-center structural neuroimaging. *Human Brain Mapping.* 2025;46:e70217. doi:10.1002/hbm.70217.

2. Martin MN, Jordanova KV, Kos AB, Russek SE, Keenan KE, Stupic KF. Relaxation measurements of an MRI system phantom at low magnetic field strengths. *Magnetic Resonance Materials in Physics, Biology and Medicine.* 2023;36:477-485. doi:10.1007/s10334-023-01086-y.
